# (In)exhaustible suppliers for evolution? Epistatic selection tunes the adaptive potential of non-genetic inheritance

**DOI:** 10.1101/294868

**Authors:** Etienne Rajon, Sylvain Charlat

**Affiliations:** Univ Lyon, Université Lyon 1, CNRS, Laboratoire de Biométrie et Biologie Evolutive UMR5558, F-69622 Villeurbanne, France

## Abstract

Non-genetic inheritance media, from methyl-accepting cytosines to culture, tend to ‘mutate’ more frequently than DNA sequences. Whether or not this makes them inexhaustible suppliers for adaptive evolution will depend on the effect of non-genetic mutations (hereafter epimutations) on fitness-related traits. Here we investigate how the magnitude of these effects might themselves evolve. More specifically, we examine the hypothesis that natural selection could set boundaries to the adaptive potential of non-genetic inheritance media due to their higher mutability. In our model, the genetic and epigenetic contributions to a non-neutral phenotype are controlled by an epistatic modifier locus, which we let evolve under the combined effects of drift and selection, in stable and in variable environments. We show that a pure genetic control evolves when the environment is stable, provided that the population is large enough, such that the phenotype becomes robust to frequent epimutations. When the environment fluctuates, however, the direction of selection on the modifier locus also fluctuates and can overall produce a large non-genetic contribution to the phenotype, especially when the epimutation rate matches the rate of environmental variation. We further show that selection on the modifier locus is mostly direct – *i.e.* it does not rely on subsequent effects in future generations – as our results are generally insensitive to recombination. These results suggest that unstable inheritance media might significantly contribute to fitness variation of traits subject to highly variable selective pressures, but little to traits responding to scarcely variable aspects of the environment, which likely represent a majority. More generally, our study demonstrates that the rate of mutation and the adaptive potential of any inheritance media should not be seen as independent properties.

---

Slowly but surely, non-genetic inheritance is making its way back into our minds (Bonduriansky 2012; Verhoeven et al. 2016). Since the discovery of DNA, biology has enjoyed a few decades of oversimplification of the much older inheritance concept, which was summarized in four letters. Yet, observations of other media of inheritance, from DNA methylation to culture, indicate that not all heritable phenotypic variation comes down to changes in DNA (Jablonka and Raz 2009; Johannes et al. 2009; Bonduriansky 2012; Danchin 2013; Quadrana and Colot 2016). How much this actually matters for evolution is the subject of a hot on-going debate (Day and Bonduriansky 2011; Laland et al. 2014; Wray et al. 2014; Charlesworth et al. 2017).

An obvious difference between genetic mutations and broadly defined epimutations (*e.g.* changes in DNA methylation patterns, histone modifications, cultural changes, etc.) is that the latter are typically more frequent (Rando and Verstrepen 2007; Johannes et al. 2009; Danchin 2013). Letting aside the heavy assumption that epimutations could also be induced by the environment (Jablonka et al. 1995; Pál 1998; Pál and Miklós 1999), the simple fact that they are frequent has led some authors to suggest that they might allow for a faster exploration of the phenotypic landscape and thus faster adaptive evolution (Klironomos et al. 2013). Subsequent models have nuanced this conclusion, pointing out that epimutations may sometimes slow down adaptation, depending on their rate of occurrence and on their effects on fitness (Furrow 2014; Kronholm and Collins 2016). While these studies incorporate the effects of these two key parameters, they ignore the possibility that they may themselves evolve and, as a result, not be independent.

Here we investigate the possibility that natural selection might adjust the fitness effects of mutations and epimutations in relation with their frequency of occurrence. In our model, we suppose that a modifier locus controls the relative genetic and epigenetic contributions to a non-neutral phenotype, in line with the observation that the effect of epimutations can be modulated by the background genome through gene / epigene epistatic interactions (Lehner 2013; Park and Lehner 2014; Blevins et al. 2017). Although mutation and epimutation rates can in principle evolve (as investigated in theoretical studies considered in more details in the Discussion) we assume here that they are fixed, to focus on their consequences on the evolution of the modifier locus.

Our results indicate that selection can effectively tune, up or down, the relative contribution of genetic and epigenetic mutations to the phenotype. In a stable environment – that is, one where the optimal phenotype remains constant across generations – a pure genetic control of the phenotype is selected for, provided that the population is large enough. In variable environments, a large epigenetic contribution is selected for immediately after an environmental change, but the opposite happens during periods of environmental stasis. Such oscillations can overall lead to large non-genetic contributions to the phenotype in variable environments, especially when the rate of epimutations matches that of environmental changes.

Our model also reveals that recombination between the trait-contributing locus (hereafter, the “trait locus”) and the modifier has little impact on the outcome. This indicates that mutations of the modifier are mainly selected through their immediate effects on their carrier, that is, through direct selection. Only in a restricted range of conditions are mutations of the modifier also affected by their interaction with future epimutations at the trait locus, that is, by indirect selection that requires tight genetic linkage with the trait locus. In that respect, the evolutionary dynamics of the “epigenetic contribution modifier” studied here contrasts markedly with that of “mutation rates modifier”, where the major role of indirect selection is well established (Ishii et al. 1989; Tenaillon et al. 2001; Salathé et al. 2009; Raynes et al. 2018). Overall, we conclude that in a wide range of conditions, the assumption of mutations and epimutations with similar phenotypic effects does not hold: depending on the context, epimutations may have greater or lower phenotypic effects than expected in the absence of selection.

## The model

We simulate the evolution of populations where non-neutral phenotypic variation is determined by a combination of two inheritance media (hereafter denoted genetic and epigenetic media, for the sake of simplicity): one with a low mutation rate *μ*_*g*_ (*g* standing for genetic) and one with a higher mutation rate *μ*_*e*_ (epigenetic). We assume that genetic and epigenetic variations affect a single locus (hereafter denoted the ‘trait locus’), so that the genetic and epigenetic states cannot be dissociated by recombination. The contribution of the two media to the phenotype depends on a ‘modifier’ locus determining *α*, the relative contribution of the epigenetic medium. Recombination between the trait and modifier loci can occur at rate *r*. We focus on the evolution of *α* under different patterns of environmental stability across generations.

A diagram of the model is presented in figure 1. Each individual i in the population is characterized by its genetic and epigenetic alleles at the trait locus (*g*_*i*_ and *e*_*i*_, respectively, taking the discrete values 0 or 1), and by its allele at the modifier locus, which sets the epigenetic contribution *α*_*i*_ varying continuously between 0 and 1. Notably, the genetic and epigenetic states at the trait locus vary independently, meaning that the epigenetic variation is ‘pure’ sensu Richards (2006). The phenotype *ϕ*_*i*_ of individual *i* is calculated as

**Figure 1.**
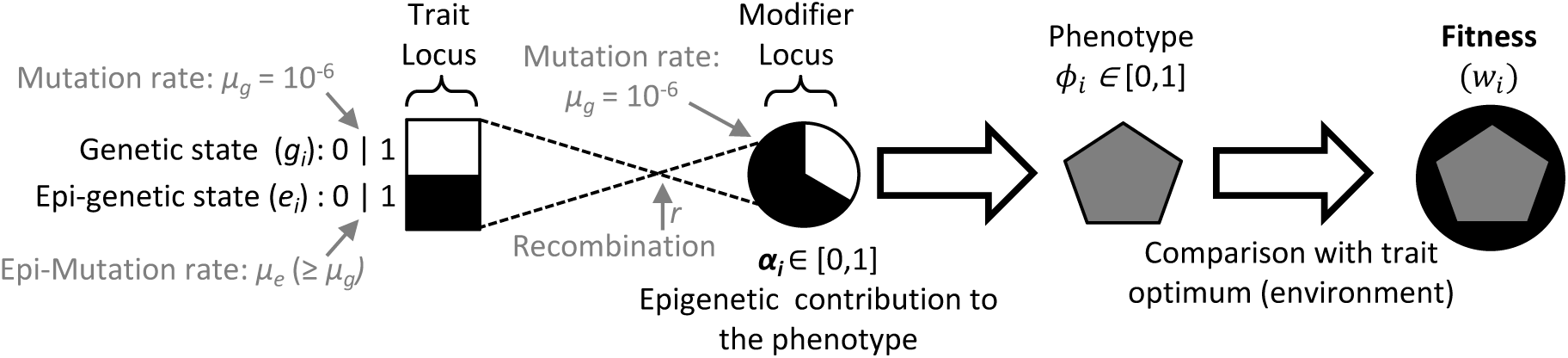
A diagram of our model. The contribution of the genetic and epigenetic states at the trait locus are modulated by the modifier locus, determining *α*, to produce the phenotype *ϕ,* which is compared to an optimum to derive fitness. The epigenetic contribution to the phenotype *α* can itself mutate with probability *μ*_*g*_. Model parameters are in grey.

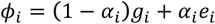

In other words, we assume that the phenotype is fully determined by the genetic and epigenetic sates, thus ignoring environmental noise that would overall reduce the heritability of the trait but would not affect the relative genetic and epigenetic contributions.

The fitness of each individual *i* at time *t, w*_*i*_(*t*), depends on the distance between its phenotype and an optimum *o*(*t*) ∈ {0,1}:

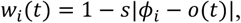

where *s* sets the strength of selection on the phenotype. The optimum can change at each generation with probability *p*_*c*_, to simulate a constant (*p*_*c*_ = 0 or varying environment (*p*_*c*_ > 0).

Populations of *N* haploid individuals (*N* = 10^4^ unless otherwise stated) are initially monomorphic with *α*_i_ sampled from a uniform distribution between 0 and 1 and *g*_*i*_ = *e*_*i*_ = *o*(0); that is, both inheritance media start at the optimal state. We then simulate their evolution through a Wright-Fisher process over 20 million generations, which we consider sufficient to reach equilibrium because results are then nearly identical to those obtained after 10 million generations (figure S1). At each generation, the population is renewed by randomly sampling the parent of each individual. The probability *p*_*i*_ that individual *i* (at *t*) is chosen as the parent of each newborn is proportional to its fitness, that is:

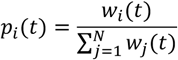

Whenever individual *i* is picked as a parent, its offspring inherit its values of *g*_*i*_, *e*_*i*_ and *α*_*i*_, unless they are modified by mutations or epimutations. The genetic mutation rate, *μ*_*g*_, equals 10^−6^ whereas the epimutation rate *μ*_*e*_ takes values from 10^−6^ to 10^−1^. The epigenetic contribution to the phenotype, *α*_*i*_, is the main variable of interest, and mutates with rate *μ*_*g*_ – i.e. we assume that *α*_*i*_ is genetically determined. Mutations of offspring *k* change *α*_*k*_(with regard to its parent’s *α*_*i*_) by an amount sampled from a normal distribution with mean 0 and standard deviation 0.1. When *α*_*k*_ decreases below 0 or increases above 1, we set it to 0 or 1, respectively. We model recombination by sampling pairs of individuals – the number of pairs is sampled from a binomial with parameters *r* and N/2 – in the new generation, and exchanging the value of *α*_*i*_ within the pair. Notably, the only difference in our model between the two inheritance media is their mutation rate; this means the model could apply in principle to situations where the trait is only determined genetically, but by two linked genes with different mutation rates.

## Results

### Evolution in a stable environment

We used simulations to investigate the evolution of the relative contribution *α* of an unstable (epigenetic) medium to a phenotypic trait under selection. We first consider a stable environment, by setting *p*_*c*_– the probability of environmental change per generation – to 0, such that the optimal phenotype does not change through time. At this stage, we also assume no recombination (*r* = 0) between the trait locus and the modifier locus controlling the value of *α*. As a preliminary control, we verified that *α* evolves neutrally when the genetic and epigenetic mutations rates are equal (*μ*_*g*_ = *μ*_*e*_ = 10^−6^; figure 2a). As expected, the distribution of the mean non-genetic contribution (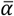) across such simulations has a mean close to 0.5 and is uniform, except that very high and low values of 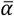 are slightly overrepresented, possibly because mutations away from the boundary states are less probable (see model section).

**Figure 2.**
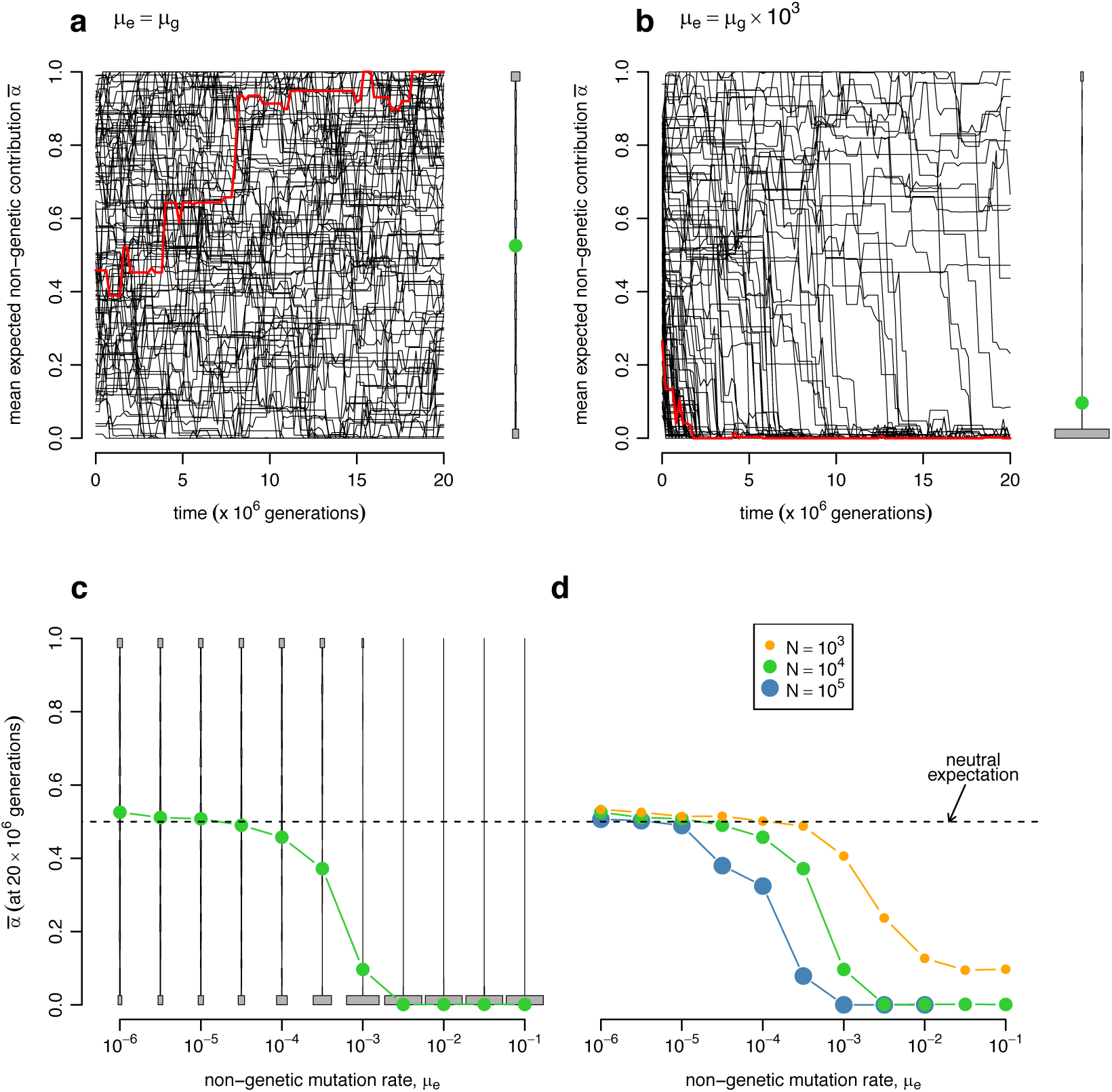
Evolutionary dynamics of 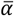 (the population average non-genetic contribution to the phenotype), in a stable environment. (a) and (b): 100 evolutionary dynamics are represented in black, with one identified in red; the final distribution of 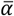 is represented on the right, summarizing 500 runs, with the mean as a green dot. This summary distribution is not informative with regard to the within-population distribution of *α*; however, we have verified it is generally unimodal (not shown). (a) When *μ*_*e*_ = *μ*_*g*_ 10^−6^ (*i.e.* when the mutation rates of both media are equal), 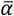 evolves neutrally in the range [0,1]. (b) When *μ*_*e*_ ≫ *μ*_*g*_ (here *μ*_*e*_ = 10^−3^ and *μ*_*g*_ = 10^−6^), 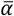 evolves toward 0 in most simulations. (c) Distributions of final values of 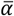 for different values of *μ*_*e*_ (with *μ*_*g*_ *=* 10^−6^) show that 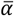 evolves away from the neutral expectation toward 0 when *μ*_*e*_ overcomes a threshold, between 10^−3.5^ and 10^−3^ in this example. (d) Mean equilibrium 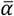 for various combinations of *μ*_*e*_ and *N*. 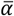 decreases when *μ*_*e*_ is above a threshold that decreases when the population is larger. Parameter values: *N* = 10^4^ (a-c), *s =* 0.01, *p*_*c*_ = 0.

In contrast, when epimutations are 1000 times more frequent than mutations (*μ*_*g*_ = 10^−6^ and *μ*_*e*_ = 10^−3^ 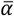 evolves to its minimal possible value in most simulations (figure 2b). This result validates the intuition that in a perfectly stable environment, selection can make the phenotype robust to frequent epimutations, that is, insensitive to their effects. A more complete picture of this process is provided in figure 2c, where we show the equilibrium distributions of 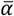 for various values of *μ*_*e*_. These simulations confirm that selection for robustness promotes a small epigenetic contribution to the phenotype, but only when *μ*_*e*_ is much larger than *μ*_*g*_, by at least 3 orders of magnitude in the context considered. When *μ*_*e*_ is below this threshold, the mean equilibrium 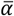 remains close to the neutral expectation of 0.5, meaning that the genetic and epigenetic media contribute equally to the phenotype.

To interpret this result, we hypothesized that selection on *α* may be too weak to oppose drift when the mutation and epimutation rates, *μ*_*g*_ and *μ*_*e*_, are close to each other. To assess the validity of this interpretation, we varied the population size and thereby the efficiency of selection. As expected, 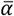 goes to 0 for smaller values of *μ*_*e*_ in larger populations, where selection is more efficient (fig 2d). Likewise, making selection more efficient through an increase of the effect of mutations on *α* (figure S2) has a similar effect. Thus, in stable environments, selection always tends to reduce the epigenetic contribution to the phenotype, but it is only effective when the population is large enough; that is, when it exceeds a threshold that depends on the epimutation rate.

We then investigated the effect of recombination between the trait and modifier loci. Figure 3 shows the evolutionarily expected value of 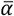 as a function of the recombination rate for different epigenetic mutation rates. Unsurprisingly, when the genetic and epigenetic mutations rates are close to each other (*e.g. μ*_*e*_ = 10^−5^, figure 3) selection on *α* remains inefficient in the presence of recombination, as it was under complete linkage. With *μ*_*e*_ = 10^−3^, recombination substantially affects the evolution of *α*, with 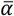 shifting from 0.1 to 0.3 for positive recombination rates – even as small as *r* ≈ 0.03. This result indicates that in the absence of recombination and for such values of *μ*_*e*_, indirect selection contributes to the evolution of *α*: alleles conferring small values of *α* carry a selective advantage because they reduce the phenotypic consequences of maladaptive epimutations at the trait locus in descendants of their carrier, a process which is only effective as long as the trait and modifier loci remain in linkage disequilibrium. Nonetheless, even with recombination, the mean 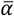 is not equal to the neutral expectation of 0.5, indicating that direct selection is also at play: mutations toward low *α* values readily confer a selective advantage because they often occur in individuals already carrying maladaptive epi-alleles, whose effect they buffer. When *μ*_*e*_ is very large (*e.g. α*_*e*_ = 10^−1^ in figure 3), we observed that recombination does not affect the equilibrium *α* values. Whatever the rate of recombination between the trait and modifier loci, a very low epigenetic contribution to the phenotype is always selected for in this context. This indicates that direct selection is here sufficient to drive the evolution of *α* toward 0.

**Figure 3.**
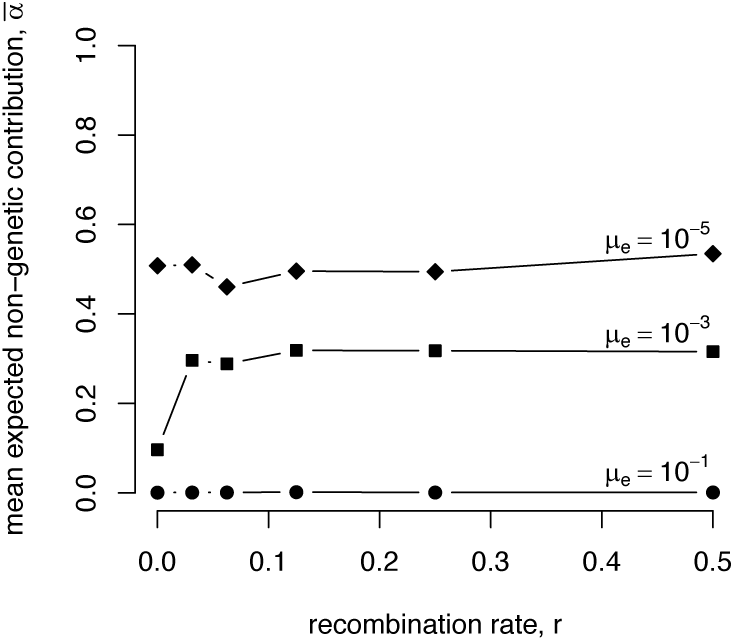
Evolutionary dynamics of the mean 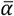 in a stable environment, with various recombination rates *r*. Parameter values: *N =* 10^4^, *s =* 0.01, **p*_*c*_ =* 0.

### Evolution in variable environments

In order to study the evolution of *α* in a non-static environment, we set the probability of environmental change *p*_*c*_ to positive values and first assume no recombination between the trait and modifier loci. The results, summarized in figure 4, indicate that the rate of environmental change has important consequences on the equilibrium value of 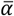. In scarcely changing environments (*e.g. p*_*c*_ = 10^−5^ in fig. 4a), high values of 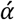 can evolve because, upon environmental change, the epigenetic medium is more often in the adaptive state than the genetic medium. However, when *μ*_*e*_ exceeds some threshold (10^−3^ for *p*_*c*_ = 10^−5^), this effect is counter-balanced by selection for robustness occurring during periods of environmental stasis. Thus, when *μ*_*e*_ is high, the evolutionarily expected value of 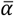 equals 0 in both stable and scarcely variable environments. In summary, selection for a high *α* does occur upon environmental changes but can be counter-balanced by selection for robustness during periods of stasis.

**Figure 4.**
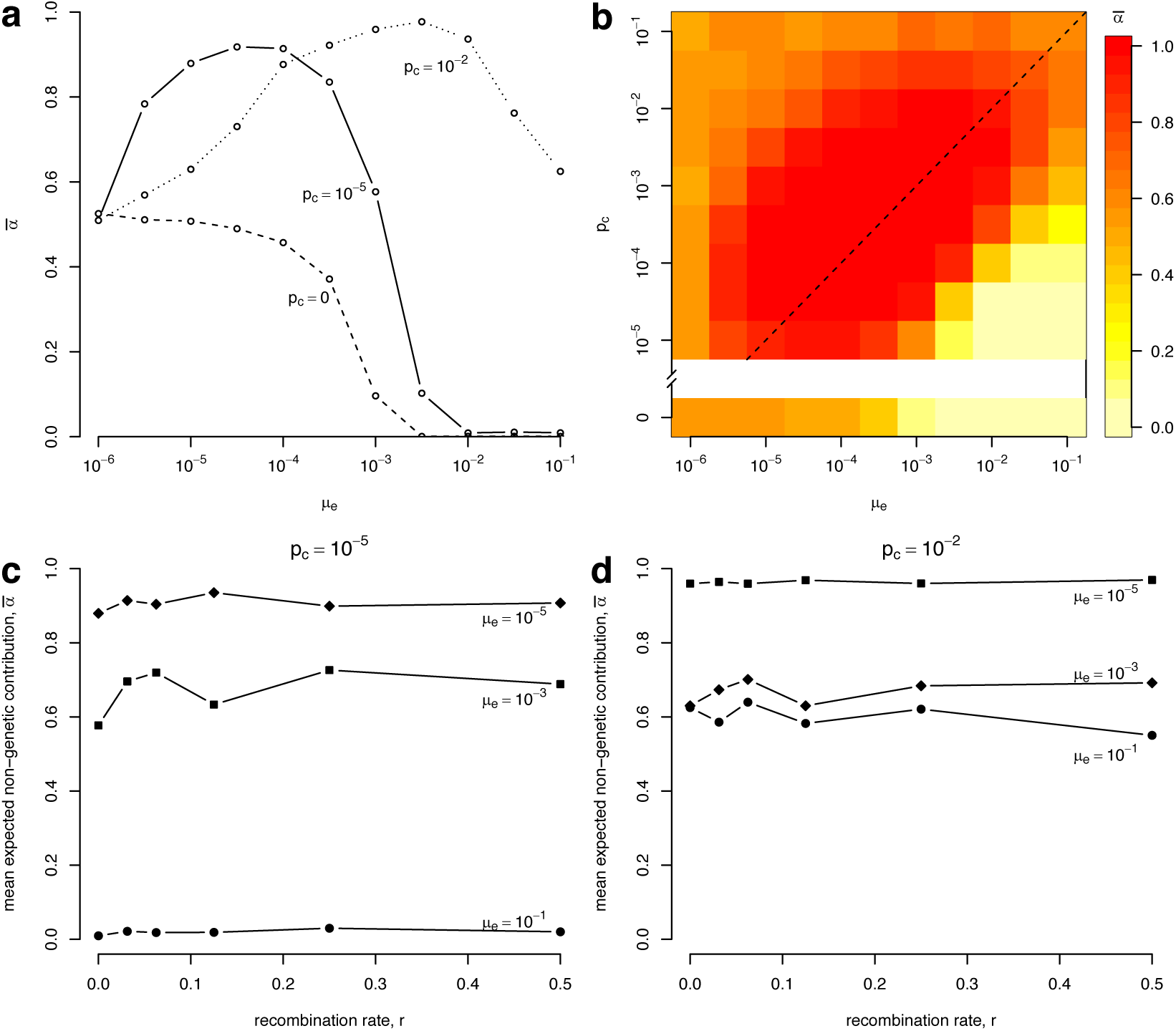
Evolutionary dynamics of 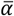 in a variable environment. (a) When the probability of environmental change *p*_*c*_ is small (10^−5^) 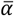 evolves above 0.5 when *μ*_*e*_ is between *μ*_*g*_ = 10^−6^ and some threshold value close to 10^−3^. In a more variable environment (*p*_*c*_ = 10^−2^ 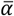 is consistently above the neutral expectation of 0.5, and peaks for higher values of *μ*_*e*_ than in the scarcely variable environment (10^−2.5^ *vs* 10^−4.5^). (b) More variable non-genetic media contribute more to the phenotype when the environment is more variable. The color scale indicates the expected values of 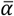. The dotted line indicates situations where *μ*_*e*_ = **p*_*c*_.* (c and d) Under the two environmental variation regimes considered in panel (a), the recombination rate has little influence on the expected value of 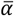. Parameter values: *N* = 10^4^, *s* = 0.01.

In line with this interpretation, increasing the probability of environmental change *p*_*c*_ (**e.g.** to 10^−2^) increases the strength of selection for large *α* values, such that selection for robustness fails to drive its evolution below the neutral expectation of 0.5, even for very high epimutation rates *μ*_*e*_ (fig 4a). Looking at a wider range of *p*_*c*_ values (fig. 4b), it appears that selection for a large contribution of the epigenetic medium is maximum when its rate of variation *μ*_*e*_ matches the frequency of environmental change (i.e. along the interrupted line in fig. 4b).

As previously illustrated in a static environment, recombination allows us to tease apart the effects of direct and indirect selection. Figures 4c and 4d show the observed mean 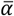 values as a function of recombination rates, for different epimutation rates and different levels of environmental instability. In all situations explored, the effect of recombination on the evolution of *α* appears to be minor, indicating that indirect selection contributes negligibly. Only in scarcely changing environments (Figure 4c) and with intermediate epimutation rates (*μ*_*e*_ = 10^−3^) does the removal of indirect selection through recombination affect the evolutionarily expected 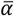, and only slightly so. This effect mirrors the trend observed in a stable environment with intermediate epimutation rates (Figure 3, *μ*_*e*_ = 10^−3^), which suggests that indirect selection contributes to drive *α* toward small values during periods of environmental stasis. Generally speaking, the very limited effect of recombination on the evolution of *α* (figure S3) suggests that direct selection largely dominates in unstable environments. In other words, depending on the rate of environmental change and epimutation rates, the epigenetic contribution to the phenotype is effectively tuned by selection, without requiring genetic linkage between the modifier and trait loci.

## Discussion

We have used simulations accounting for the effect of drift and selection to unravel the relationship between the instability of a non-genetic inheritance medium – its mutation rate – and its evolving contribution to a non-neutral phenotype, termed *α* in our model. Before any attempt to draw general conclusions, we should stress that our analysis is based on an abstract model, which is unlikely to reach the level of realism required to obtain a precise expectation for this relationship for any particular inheritance medium. Instead of being specific and predictive, our aim here was to formally explore the conjecture that the adaptive potential of an inheritance media should depend on its transmission accuracy – or, inversely, on its mutation rate *sensu lato* – in interaction with the (un)stability of the environment. We believe that this approach helps clarify when non-genetic inheritance may have important evolutionary implications and brings relevant elements in the complex ongoing debate on this question.

One important and clear outcome of our analysis is that the relationship between the epimutation rate and the epigenetic contribution to the phenotype can be shaped by epistatic selection and is thus generally not flat. Differences in phenotypic effects between inheritance media will translate into different adaptive potentials, even though the relationship between mutational size effect and adaptive potential can be complex and possibly non monotonic (Fisher 1930; Kronholm and Collins 2016). The link between mutation rates and phenotypic effects indicates that the assumption of identical contributions to the phenotype of genetic and non-genetic media, which has led some authors to conclude that non-genetic inheritance can lead to rapid adaptive evolution (Klironomos et al. 2013), is not tenable in principle. Specifically, in scarcely varying environments, selection for robustness is expected to reduce the phenotypic contribution of any medium with a high mutation rate (figure 2). However, for a given population size, this selective process is only effective when the mutation rate is above a threshold. Reciprocally, the phenotypic contribution of an inheritance media with a given mutation rate will either evolve neutrally or be under efficient selection, depending on the population size.

We have also found that environmental instability can favor the evolution of large non genetic contributions to the phenotype, especially when the epimutation rate matches the rate of environmental variation, as anticipated by Danchin (2013). This conclusion can be interpreted in the light of previous theoretical work in various areas of evolutionary biology, focusing for instance on the evolution of mutation rates (Ishii et al. 1989; Salathé et al. 2009), of the rates of switching between phenotypic states (Lachmann and Jablonka 1996; Thattai and Van Oudenaarden 2004; Kussell and Leibler 2005; Wolf et al. 2005; King and Masel 2007; Rando and Verstrepen 2007; Gaal et al. 2010; Liberman et al. 2011; Mayer et al. 2017) and of bet-hedging strategies (Cohen 1966; Seger and Brockmann 1987; Clauss and Venable 2000; Rajon et al. 2009, 2014; Carja and Plotkin 2017). This body of work generally predicts that fitness is maximized when the switching rate (that is, the mutation rate in a broad sense) matches the rate of environmental change. In our model, this situation would correspond to *μ*_*e*_ ≈ *p*_*c*_ combined with a maximal contribution of the non-genetic media (*α* = 1). Departing from this ideal situation by decreasing *α* would necessarily decrease fitness, because the phenotype would then rely on both an optimally changing (non-genetic) and a sub-optimally changing (genetic) media. Accordingly, *α* evolves in our model such that the contribution of the non-genetic media is maximum when *α*_*e*_ and *p*_*c*_ are close; as they differ, a joint contribution of both media (*α* < 1) evolves.

Despite the apparent similarity between the aforementioned body of work and the present study, the effect of recombination between the modifier and trait loci reveals an important difference. Indeed, our model shows that both direct and indirect selection can impact the evolution of the epigenetic contribution to the phenotype, with a relative weight that depends on the epimutation rate. In fact, in most situations direct selection is sufficient to explain the evolution of this contribution. This means that even though they act through epistatic interactions, mutations modifying the epigenetic contribution can readily gain a selective advantage (direct selection) if they land on a genetic background where buffering or enhancing the effect of an epimutation improves fitness. This is in sharp contrast with current views on the evolution of mutation rates: these are only subject to indirect selection, which requires strong linkage, because their potential selective advantage only relies on the future mutations they will or will not generate.

The probability of environmental change *p*_*c*_ plays a major role in our results, as the occurrence of an environmental change gives a selective advantage to genotypes with an increased epigenetic contribution *α*. It is interesting to realize that different patterns of environmental stability could correspond, not to different environments *per se*, but to different traits, each related to more or less stable aspects of the environment. For instance, traits responding to gravity should be under extremely stable selection, whereas other traits – **e.g.** those involved in immunity or the exploitation of resources whose abundances vary in time – should be under ever-changing selection pressures. Our analysis suggests that different inheritance media – characterized by different mutation rates – should have different relative contributions to these different categories of traits.

We should also clarify that, even though our graphical representation in fig. 4b seems to indicate that large values of *α* are expected in most situations, our choice of what situations are modeled is arbitrary and unlikely representative of reality. For instance, there is no reason to assume that traits facing highly unstable environments (*e.g.* with *p*_*c*_ = 10^−2^) are as common as those facing stable environments (with *p*_*c*_ = 0). On the contrary, many pieces of evidence together argue that stabilizing selection is largely prevalent in nature. It is now well established from paleontological records that evolutionary stasis is often observed at the morphological level over long periods, which seems best explained by long episodes of stabilizing selection (Estes and Arnold 2007). This macro-evolutionary pattern is compatible with the general picture drawn by micro-evolutionary studies aimed at describing selection in contemporary populations. Indeed, meta-analyses of such studies have concluded that strong directional selection is rare (Kingsolver et al. 2001; Morrissey and Hadfield 2012; Arnold 2014). Likewise, at the molecular level, the fact that the majority of non-neutral sites within genomes are under purifying selection advocates for the idea that most traits are related to stable aspects of the environment (Eyre-Walker and Keightley 2007).

Notably, we have assumed that any mutation or epimutation deleterious at time *t* may become beneficial in the future if the environment changes. In other words, contrary to other population genetics models (Johnson 1999; Rajon and Masel 2011) we have ignored unconditionally deleterious mutations and epimutations. The presence of such mutations might increase the strength of selection for robustness and, overall, make the evolution of large epigenetic contributions less likely than described here. Whether unconditionally deleterious epimutations do exist, however, is unclear. In fact, since most deleterious mutations prevent protein folding (Wylie and Shakhnovich 2011) – which non genetic mutations can hardly do – epimutations may be less likely deleterious than mutations. The distribution of their fitness effect is, obviously, an important parameter to consider in future studies.

Even among mutations and epimutations with potentially adaptive effects, the present study ignores potential differences in the *qualitative* effects of mutations and epimutations, hence focusing on situations where the genetic and non-genetic media impact similar aspects of the phenotype. This matches, for example, changes in gene expression that can be modulated by genetic changes in regulatory sequences and, indistinguishably, by gene methylation or histone covalent modifications. In contrast, only genetic mutations can change a protein sequence, with almost endless possible phenotypic consequences (Kronholm and Collins 2016). The information content of an inheritance medium has been suggested to set an upward limit to its mutation rate (Eigen and Schuster 1977; Maynard Smith 1990), which might explain why media with more possible states evolve lower mutation rates. Our work suggests that this relationship might work both ways, such that the mutation rate of an inheritance medium also impacts its evolutionary potential. This hypothesis could be addressed by modeling the evolution of the phenotypic space that a medium can access, instead of (or in addition to) its relative contribution to the phenotype as considered in this study. We anticipate that within such a framework, the most variable (non-genetic) medium should evolve a mutational space that only includes frequently occurring phenotypic optima, excluding other mutational targets due to selection for robustness. The least variable (genetic) medium should instead keep access to a wider set of phenotypes due to the comparatively weaker selection for robustness. Interestingly, if and when this prediction holds, the non-genetic medium may still produce higher phenotypic variance than the genetic medium due to its higher mutation rate, and therefore contribute more to heritability. This indicates that the contribution of one medium to heritability potentially ignores the diversity of phenotypes it can access. Because the potential for future evolutionary innovation certainly depends on this diversity, the link between a medium’s contribution to heritability and its long-term adaptive potential may well be tenuous.

## Conclusion

The present study was aimed at providing some consideration to the evolutionary implication of non-genetic inheritance. While the question is obviously not settled, we would argue that at this stage, acknowledging that non-genetic inheritance *might* contribute to adaptive evolution is already a big step forward. The power of the genetic paradigm is not in question: a huge variety of studies have identified clear causal links between genetic variation, phenotypes and selective pressures, demonstrating that genetic evolution can explain major evolutionary changes (*e.g.* Lenormand et al. 1999; Levy and Marshall 2004; Jones et al. 2012). Similarly, developmental genetics have provided ample evidence that important macro-evolutionary changes come down to genetics (Stern and Orgogozo 2009). However, it is notable that the huge part of evolution explained by genetics tells us little about the potentially also huge part that is not explained by genetics. The strength of evolutionary genetics largely holds in the tractability of genetic changes: genomes can be sequenced, aligned, and substitutions can be identified. On the contrary, non-genetic inheritance encompasses a wide variety of processes, which are hardly tractable. The present study has focused on one particular parameter: the rate of random change from one generation to the next. Although this is obviously an insufficient proxy to catch the subtleties of all inheritance processes, our results indicate that it is one that sets important constraints to their evolutionary potential.

---

**Figure S1.**
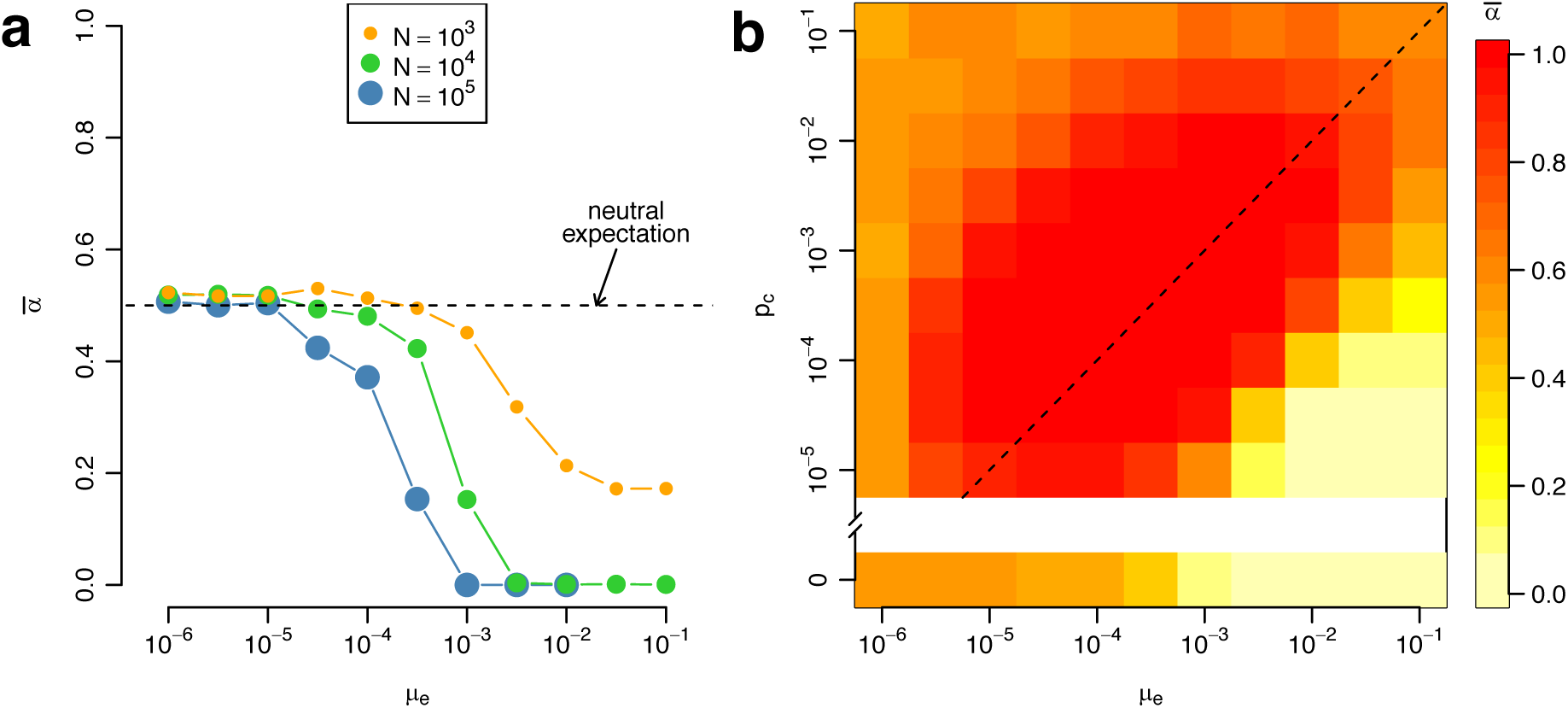
Evolutionary expectations of 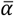 after 10 million generations are almost identical to those obtained with our default simulation time of 20 million generations. Compare panel (a) to figure 2d, and panel (b) to figure 3b. Aside from simulation time, parameters are identical to those described in the legends of these figures. Notably, the 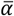 value reached after 10 million generations in the smallest population (N=10^3^) is higher than after 20 million generations (figure 2d) indicating that its evolution is very slow in this context due to ineffective selection.

**Figure S2.**
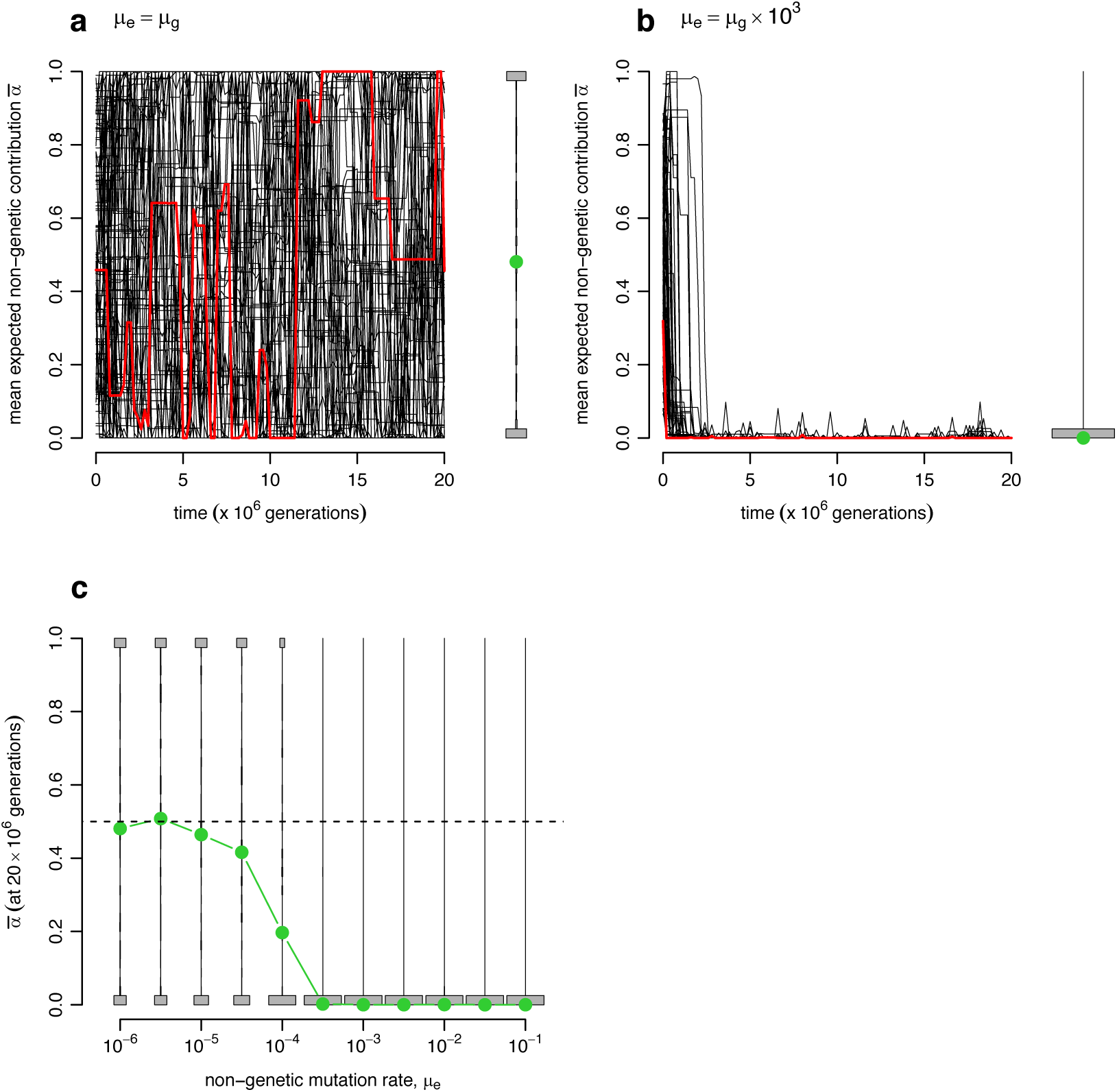
The equivalent of panels (a), (b) and (c) in figure 2, but with larger effects of mutations on *a* (the standard deviation of the distribution of mutational effects equals 0.5 instead of 0.1). In such conditions, selection on *α* is stronger and thus effective for lower values of *μ*_*e*_.

**Figure S3.**
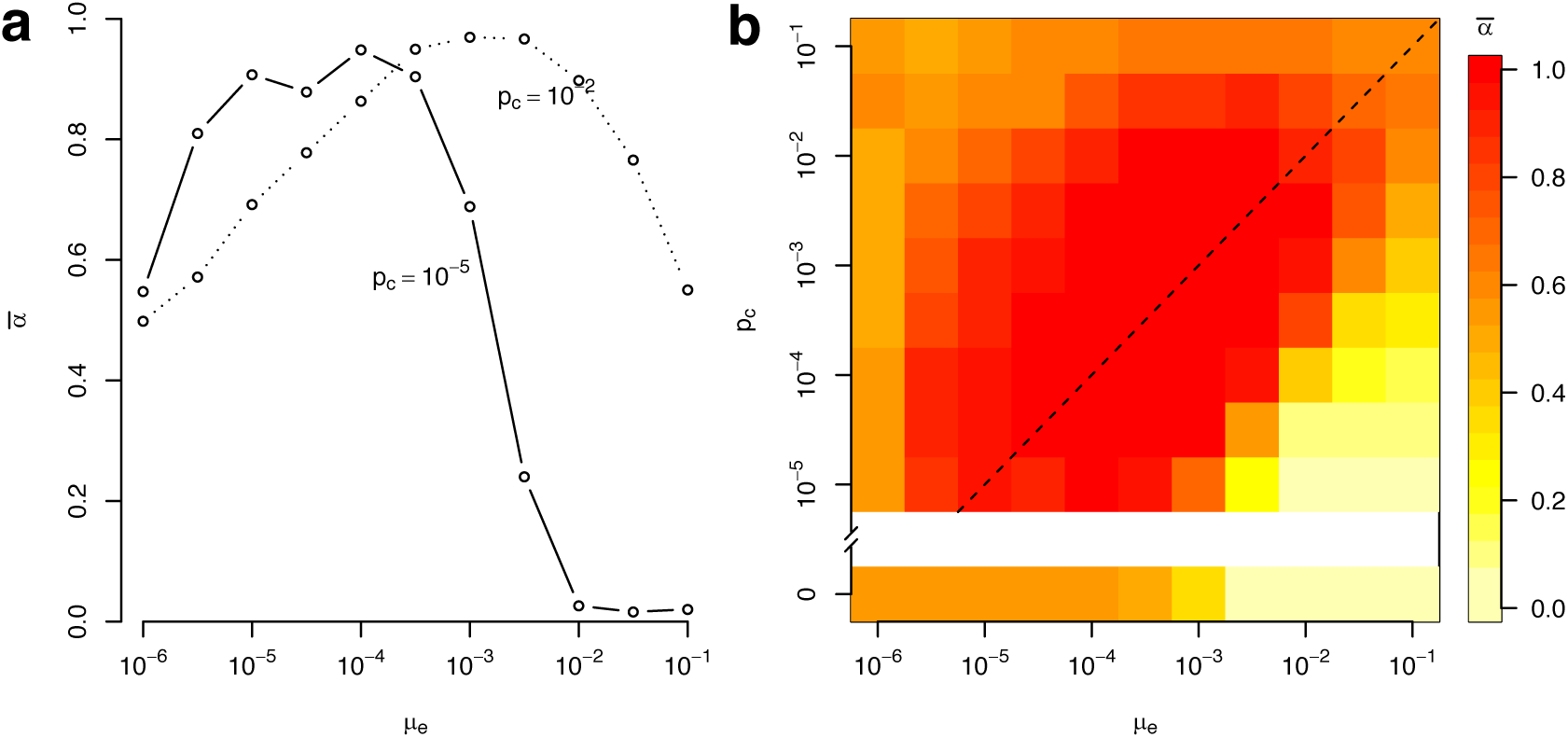
The equivalent, with recombination (*r*=0.5), of panels (a) and (b) in figure 3.

## References

Arnold, S. J. 2014. Phenotypic Evolution: The Ongoing Synthesis. The American Naturalist 183: 729–746.

Blevins, T., J. Wang, D. Pflieger, F. Pontvianne, and C. S. Pikaard. 2017. Hybrid incompatibility caused by an epiallele. Proceedings of the National Academy of Sciences 114: 3702–3707.

Bonduriansky, R. 2012. Rethinking heredity, again. Trends in Ecology and Evolution 27: 330–336.

Carja, O., and J. B. Plotkin. 2017. The evolutionary advantage of heritable phenotypic heterogeneity. Scientific Reports 7: 1–12.

Charlesworth, D., N. H. Barton, and B. Charlesworth. 2017. The sources of adaptive variation. Proceedings of the Royal Society B: Biological Sciences 284: 20162864.

Clauss, M., and D. L. Venable. 2000. Seed Germination in Desert Annuals: An Empirical Test of Adaptive Bet Hedging. American Naturalist 155: 168–186.

Cohen, D. 1966. Optimizing reproduction in a randomly varying environment. Journal of Theoretical Biology 12: 119–129.

Danchin, E. 2013. Avatars of information: Towards an inclusive evolutionary synthesis. Trends in Ecology and Evolution 28: 351–358.

Day, T., and R. Bonduriansky. 2011. A Unified Approach to the Evolutionary Consequences of Genetic and Nongenetic Inheritance. The American Naturalist 178:E18–E36.

Eigen, M., and P. Schuster. 1977. The Hypercyde. Die Naturwissenschaften 64: 541.

Estes, S., and S. J. Arnold. 2007. Resolving the Paradox of Stasis: Models with Stabilizing Selection Explain Evolutionary Divergence on All Timescales. The American Naturalist 169: 227–244.

Eyre-Walker, A., and P. D. Keightley. 2007. The distribution of fitness effects of new mutations. Nature reviews. Genetics 8: 610–8.

Fisher, R. A. 1930. The genetical theory of natural selection. Oxford University Press, Oxford.

Furrow, R. E. 2014. Epigenetic inheritance, epimutation, and the response to selection. PLoS ONE 9: 7–10.

Gaal, B., J. W. Pitchford, and A. J. Wood. 2010. Exact results for the evolution of stochastic switching in variable asymmetric environments. Genetics 184: 1113–1119.

Ishii, K., H. Matsuda, Y. Iwasa, and A. Sasaki. 1989. Evolutionarily stable mutation rate in a periodically changing environment. Genetics 121: 163–174.

Jablonka, E., B. Oborny, I. Molnár, E. Kisdi, J. Hobfauer, and T. Czárán. 1995. The adaptive advantage of phenotypic memory in changing environments 350: 133–141.

Jablonka, E., and G. Raz. 2009. Transgenerational Epigenetic Inheritance: Prevalence, Mechanisms, and Implications for the Study of Heredity and Evolution. The Quarterly Review of Biology 84: 131–176.

Johannes, F., E. Porcher, F. K. Teixeira, V. Saliba-Colombani, M. Simon, N. Agier, A. Bulski, et al. 2009. Assessing the impact of transgenerational epigenetic variation on complex traits. PLoS Genetics 5.

Johnson, T. 1999. Beneficial mutations, hitchhiking and the evolution of mutation rates in sexual populations. Genetics 151: 1621–1631.

Jones, F. C., M. G. Grabherr, Y. F. Chan, P. Russell, E. Mauceli, J. Johnson, R. Swofford, et al. 2012. The genomic basis of adaptive evolution in threespine sticklebacks. Nature 484: 55–61.

King, O., and J. Masel. 2007. The evolution of bet-hedging adaptations to rare scenarios. Theoretical Population Biology 72: 560–575.

Kingsolver, J. G., H. E. Hoekstra, J. M. Hoekstra, D. Berrigan, S. N. Vignieri, C. E. Hill, A. Hoang, et al. 2001. The Strength of Phenotypic Selection in Natural Populations. The American Naturalist 157: 245–261.

Klironomos, F. D., J. Berg, and S. Collins. 2013. How epigenetic mutations can affect genetic evolution: Model and mechanism. BioEssays 35: 571–578.

Kronholm, I., and S. Collins. 2016. Epigenetic mutations can both help and hinder adaptive evolution. Molecular Ecology 25: 1856–1868.

Kussell, E., and S. Leibler. 2005. Phenotypic Diversity, Population Growth, and Information in Fluctuating Environments. Science 309: 2075.

Lachmann, M., and E. Jablonka. 1996. The inheritance of phenotypes: an adaptation to fluctuating environments. Journal of theoretical biology 181: 1–9.

Laland, K., T. Uller, M. Feldman, K. Sterelny, G. B. Müller, A. Moczek, E. Jablonka, et al. 2014. Does Evolutionary Theory Need A Rethink? Yes, urgently. Nature 514: 161–164.

Lehner, B. 2013. Genotype to phenotype: lessons from model organisms for human genetics. Nature reviews. Genetics 14: 168–78.

Lenormand, T., D. Bourguet, T. Guillemaud, and M. Raymond. 1999. Tracking the evolution of insecticide resistance in the mosquito Culex pipiens. Nature 400: 861–864.

Levy, S. B., and B. Marshall. 2004. Antibacterial resistance worldwide: causes, challenges and responses. Nature Medicine 10: S122–S129.

Liberman, U., J. Van Cleve, and M. W. Feldman. 2011. On the evolution of mutation in changing environments: Recombination and phenotypic switching. Genetics 187: 837–851.

Mayer, A., T. Mora, O. Rivoire, and A. M. Walczak. 2017. Transitions in optimal adaptive strategies for populations in fluctuating environments. arXiv preprint 1–18.

Maynard Smith, J. 1990. Models of a Dual Inheritance System. Journal of theoretical biology 143: 41–53.

Morrissey, M. B., and J. D. Hadfield. 2012. Directional selection in temporally replicated studies is remarkably consistent. Evolution 66: 435–442.

Pál, C. 1998. Plasticity, memory and the adaptive landscape of the genotype. Proceedings of the Royal Society B: Biological Sciences 265: 1319–1323.

Pál, C., and I. Miklós. 1999. Epigenetic inheritance, genetic assimilation and speciation. Journal of theoretical biology 200: 19–37.

Park, S., and B. Lehner. 2014. Epigenetic epistatic interactions constrain the evolution of gene expression. Molecular Systems Biology 9: 645–645.

Quadrana, L., and V. Colot. 2016. Plant Transgenerational Epigenetics. Annual Review of Genetics 50: 467–491.

Rajon, E., E. Desouhant, M. Chevalier, F. Débias, and F. Menu. 2014. The Evolution of Bet Hedging in Response to Local Ecological Conditions. The American Naturalist 184:E1–E15.

Rajon, E., and J. Masel. 2011. Evolution of molecular error rates and the consequences for evolvability. Proceedings of the National Academy of Sciences 108: 1082–1087.

Rajon, E., S. Venner, and F. Menu. 2009. Spatially heterogeneous stochasticity and the adaptive diversification of dormancy. Journal of Evolutionary Biology 22: 2094–2103.

Rando, O. J., and K. J. Verstrepen. 2007. Timescales of Genetic and Epigenetic Inheritance. Cell 128: 655–668.

Raynes, Y., C. S. Wylie, P. D. Sniegowski, and D. M. Weinreich. 2018. Sign of selection on mutation rate modifiers depends on population size Yevgeniy. Proc Natl Acad Sci U S A in press.

Richards, E. J. 2006. Revisiting Soft Inheritance 7: 395–402.

Salathé, M., J. Van Cleve, and M. W. Feldman. 2009. Evolution of stochastic switching rates in asymmetric fitness landscapes. Genetics 182: 1159–1164.

Seger, J., and H. J. Brockmann. 1987. What is bet-hedging? Pages 182–211 in P. Harvey and L. Partridge, eds. Oxford Surveys in Evolutionary Biology (Vol. 4). Oxford Univeristy Press.

Stern, D., and V. Orgogozo. 2009. Is Genetic Evolution Predictable? Science 323: 746–752.

Tenaillon, O., F. Taddei, M. Radman, and I. Matic. 2001. Second-order selection in bacterial evolution: Selection acting on mutation and recombination rates in the course of adaptation. Research in Microbiology 152: 11–16.

Thattai, M., and A. Van Oudenaarden. 2004. Stochastic gene expression in fluctuating environments. Genetics 167: 523–530.

Verhoeven, K. J. F., B. M. VonHoldt, and V. L. Sork. 2016. Epigenetics in ecology and evolution: What we know and what we need to know. Molecular Ecology 25: 1631–1638.

Wolf, D. M., V. V. Vazirani, and A. P. Arkin. 2005. Diversity in times of adversity: Probabilistic strategies in microbial survival games. Journal of Theoretical Biology 234: 227–253.

Wray, G. A., H. E. Hoekstra, D. J. Futuyma, R. E. Lenski, T. F. C. Mackay, D. Schluter, and J. E. Strassmann. 2014. Does evolutionary theory need a rethink? No, all is well. Nature 514: 161–164.

Wylie, C. S., and E. I. Shakhnovich. 2011. A biophysical protein folding model accounts for most mutational fitness effects in viruses. Proceedings of the National Academy of Sciences 108: 9916–9921.

